# Evaluation of Deep Neural Network ProSPr for Accurate Protein Distance Predictions on CASP14 Targets

**DOI:** 10.1101/2021.10.14.464472

**Authors:** Jacob Stern, Bryce Hedelius, Olivia Fisher, Wendy M. Billings, Dennis Della Corte

## Abstract

The field of protein structure prediction has recently been revolutionized through the introduction of deep learning. The current state-of-the-art tool AlphaFold2 can predict highly accurate structures, however, it has a prohibitively long inference time for applications that require the folding of hundreds of sequences. The prediction of protein structure annotations, such as amino acid distances, can be achieved at a higher speed with existing tools, such as the ProSPr network. Here, we report on important updates to the ProSPr network, its performance on the recent Critical Assessment of Structure Prediction (CASP14) competition, and an evaluation of its accuracy dependency on multiple sequence alignment depth. We also provide a detailed description of the architecture and the training process, accompanied by reusable code. This work is anticipated to provide a solid foundation for the further development of protein distance prediction tools.

## 1. Introduction

Proteins are among Natures’ smallest machines and fulfill a broad range of life-sustaining tasks. To fully understand the function of a protein, accurate knowledge of its folded structure is required. Protein structures can either be obtained from experiments, homology modeling, or computational structure prediction. Accurate structures can be used for the rational design of biosensors [1], prediction of small-molecule docking [2], enzyme design [3], or simulation studies to explore protein dynamics. [4]

Recent progress in the field of computational structure prediction includes end-2-end deep learning models Alphafold2 [5] and RoseTTAfold [7] that are able to predict highly accurate protein structures from multiple sequence alignments. Alphafold2 has been used to predict the structures of many protein sequences found in Nature, including the human proteome.[6] Despite these advancements, it is still not fully known if models such as Alphafold2 can extract dynamics or multiple conformations of proteins.[7] Furthermore, it is also not clear if Alphafold2 can be used effectively to support tasks in protein engineering, such as assessing if single point mutations in the amino acid sequence of a protein will alter stability or function.

A main bottleneck of Alphafold2 is the runtime for prediction. It can take multiple hours on a GPU cluster to predict the structure of a single protein. If thousands of sequences must be evaluated in a protein design study, this runtime can be prohibitive. Previous state-of-the-art tools Alphafold1[8] and trRosetta[9] used a two-stage process to predict protein folds. The first being the prediction of distances and contacts between amino acids. This task can be performed rapidly and allows for comparing difference between contact patterns of multiple sequences.

After Alphafold1 was initially presented during the Critical Assessment of Protein Structure Prediction (CASP13) conference [10], many questions remained about its implementation. To demystify this process, our team developed and published ProSPr – a clone of Alphafold1 on GitHub and bioRxiv.[11] With the release of the Alphafold1 paper, we updated the ProSPr architecture and made new models available. After CASP14, it became apparent that ProSPr was used by multiple participating groups, as the Alphafold1 code was not easily usable by the community.[14]

Deep learning methods are often complementary, and a variety of easy-to-use models can be very valuable to form ensembles that outperform single methods. In a previous study, we have shown that ProSPr contact predictions are of similar quality as Alphafold1 and trRosetta predictions but that an ensemble of all three methods is superior to any individual method.[12] We further showed that ProSPr can be used to rapidly predict large structural changes from small sequence variations, making it a useful tool for sequence assessment in protein engineering.[12]

While the first ProSPr model has been used by multiple groups during CASP14 and shown its usefulness in driving improved contact predictions, this is the first detailed description of its updated architecture and the training process used. We did not use our original version of ProSPr in CASP14, but rather a completely distinct iteration with higher performance that drew from our growing expertise in the area. These updates were informed by the publication of Alphafold1 and trRosetta that were not released until shortly before the CASP14 prediction season began, and so the models described here were still being trained during CASP14 and are distinct from those we used during the competition. Here, we present this improved ProSPr version and release the network code, training scripts, and related datasets.

## 2. Evaluation and Results

We evaluated the performance of three updated ProSPr models using the CASP14 target dataset. The CASP assessors provided access to full label information before it was publicly available (i.e., prior to PDB release) for many of the targets which enabled us to analyze our predictions across 61 protein targets. We evaluated these targets based on residue-residue contacts, which are defined by CASP as having a Cβ (or Cα for glycine) distance less than 8 Å.[13] Predicted contact probabilities were straightforward to derive from our binned distance predictions: we summed the probabilities of the first three bins since their distances correspond to those less than 8 Å.

Figure 1 shows results for two example targets from CASP14. For T1034 we were able to construct an MSA with a depth greater than 10,000 and the predicted accuracies (top of the diagonal) are in good agreement with the labels (bottom of the diagonal). The protein structure annotations on the right compare the prediction accuracy on top with the label on the bottom. This shows that even for an easy target, these predictions are not highly accurate, which is likely due to the small loss contribution assigned to auxiliary predictions (see Methods). For target T1042 no sequences could be found, and the corresponding predictions are without signal. The goal of training a contact prediction tool that can infer information from sequence alone is an open problem and will need to be addressed in future work.

**Figure 1.**
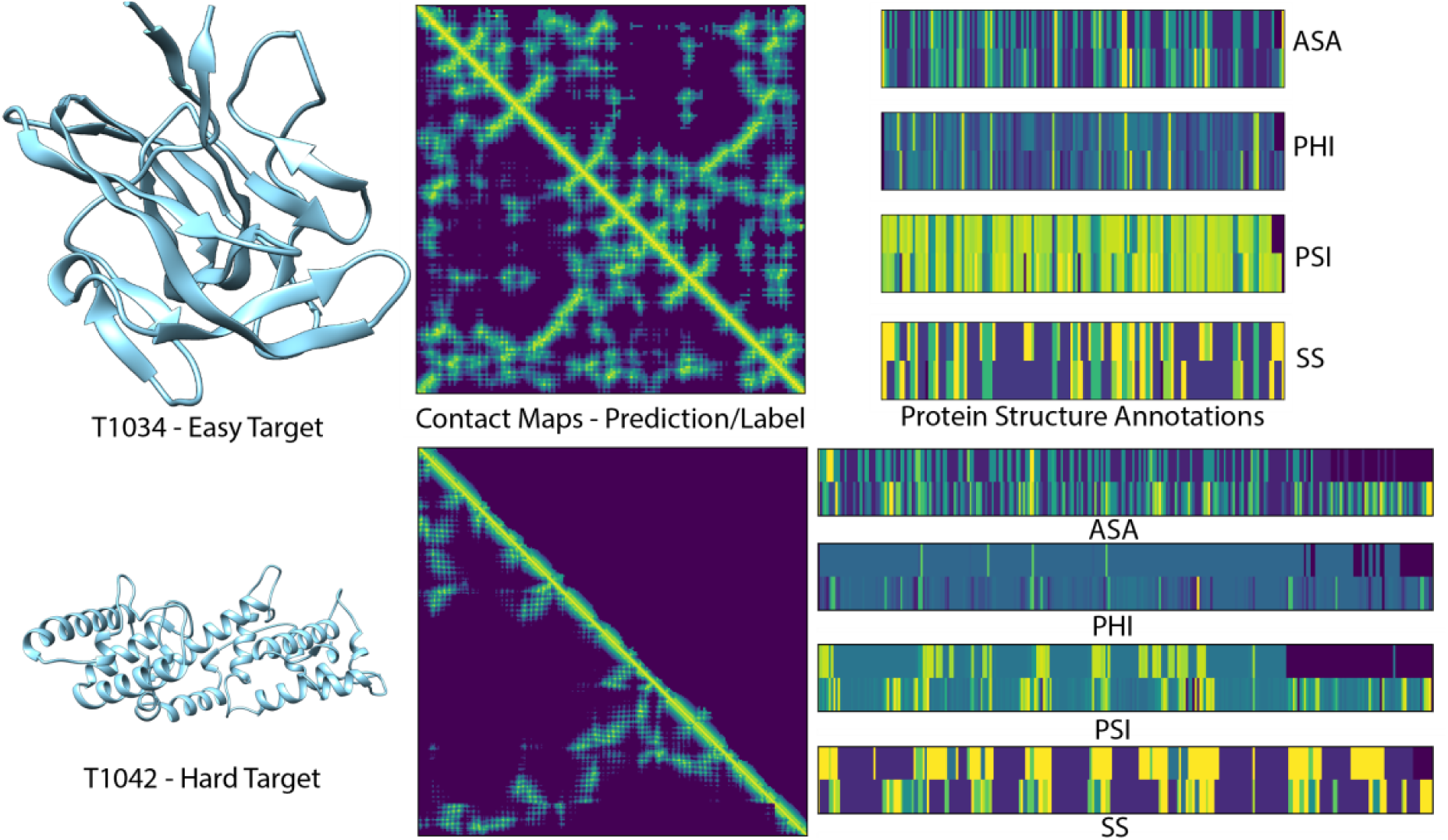
Two example targets from CASP14 test set. On left experimental structures from which labels were derived. In middle, contact maps predicted with ProSPr ensemble on top of diagonal, label on bottom. On right, visualization of auxiliary loss predictions on top with labels at bottom: Accessible Surface Area (ASA), Torsion angles (PHI, PSI), Secondary Structure (SS).

Table 1 shows the contact accuracies of the three ProSPr models evaluated at short, mid, and long contact ranges. These categories relate to the sequence separation of the two amino acids involved in each contact, where short, mid, and long-range pairs are separated by 6 to 11, 12 to 23, and 24+ residues, respectively. [14] All contact predictions in each of these ranges were ranked by probability and the top L (sequence length) pairs in each category were considered to be in contact. We then calculated contact accuracies using the following equation [15]:

**Table 1.** CASP14 Contact Accuracies.

| ProSPr Model | Contact Accuracy (%) |  |  |  |
| --- | --- | --- | --- | --- |
|  | Short | Mid | Long | Average |
| A | 81.09% | 69.52% | 41.63% | 64.08% |
| B | 81.15% | 69.29% | 42.41% | 64.28% |
| C | 81.94% | 69.97% | 43.59% | 65.17% |
| Ensemble | **82.08%** | **70.55%** | **44.04%** | **65.56%** |

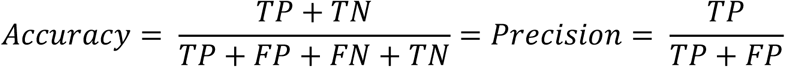

which reduces to precision since no negative predictions are made *(TN = FN = 0)*. Furthermore, we normalized the accuracy scores for each target in each range so that the full range of 0–100% could be achieved (i.e. in some cases there may not be L true contacts, so the maximum score would otherwise be lower).

The three ProSPr models shown in Table 1 have the same architecture and were trained on the same data (see Methods) but perform somewhat differently. By creating an ensemble of the three networks, the average results in all three areas are improved (for ensemble performance on individual targets, see Table 2) which is in accordance with our previous work [12]. We have made all three models individually available, but in accordance with these results, the default inference setting of the code is to automatically ensemble all of them for the best performance.

**Table 2.** ProSPr Ensemble Contact Accuracies.

| Target | Contact Accuracy |  |  |
| --- | --- | --- | --- |
|  | Short | Mid | Long |
| T1045s2 | 0.833 | 0.924 | 0.694 |
| T1046s1 | 1.000 | 1.000 | 0.536 |
| T1046s2 | 0.892 | 0.574 | 0.303 |
| T1047s1 | 0.907 | 0.985 | 0.639 |
| T1047s2 | 1.000 | 0.983 | 0.852 |
| T1060s2 | 0.857 | 0.575 | 0.282 |
| T1060s3 | 0.976 | 0.955 | 0.793 |
| T1065s1 | 1.000 | 0.973 | 0.518 |
| T1065s2 | 1.000 | 1.000 | 0.870 |
| T1024 | 1.000 | 1.000 | 0.809 |
| T1026 | 0.750 | 0.425 | 0.494 |
| T1027 | 0.485 | 0.278 | 0.054 |
| T1029 | 0.891 | 0.818 | 0.220 |
| T1030 | 0.804 | 0.792 | 0.333 |
| T1031 | 0.686 | 0.457 | 0.105 |
| T1032 | 0.889 | 0.851 | 0.580 |
| T1033 | 0.750 | 0.316 | 0.216 |
| T1034 | 0.988 | 0.874 | 0.885 |
| T1035 | 0.412 | 0.080 | 0.000 |
| T1037 | 0.690 | 0.455 | 0.030 |
| T1038 | 0.720 | 0.538 | 0.407 |
| T1039 | 0.269 | 0.000 | 0.007 |
| T1040 | 0.318 | 0.222 | 0.027 |
| T1041 | 0.644 | 0.357 | 0.021 |
| T1042 | 0.487 | 0.441 | 0.058 |
| T1043 | 0.431 | 0.216 | 0.014 |
| T1049 | 1.000 | 0.939 | 0.440 |
| T1050 | 0.964 | 0.821 | 0.705 |
| T1052 | 0.728 | 0.600 | 0.417 |
| T1053 | 0.796 | 0.521 | 0.093 |
| T1054 | 1.000 | 1.000 | 0.710 |
| T1055 | 0.932 | 0.860 | 0.200 |
| T1056 | 0.823 | 0.829 | 0.661 |
| T1057 | 1.000 | 0.987 | 0.815 |
| T1058 | 0.821 | 0.678 | 0.678 |
| T1061 | 0.807 | 0.687 | 0.511 |
| T1064 | 0.615 | 0.500 | 0.094 |
| T1067 | 0.865 | 0.824 | 0.466 |
| T1068 | 0.926 | 0.813 | 0.204 |
| T1070 | 0.941 | 0.707 | 0.579 |
| T1073 | 1.000 | 1.000 | 1.000 |
| T1074 | 0.845 | 0.700 | 0.328 |
| T1076 | 0.970 | 0.947 | 0.911 |
| T1078 | 0.984 | 0.892 | 0.587 |
| T1079 | 0.956 | 0.964 | 0.739 |
| T1082 | 0.615 | 0.636 | 0.164 |
| T1083 | 0.909 | 0.783 | 0.909 |
| T1084 | 1.000 | 1.000 | 1.000 |
| T1087 | 1.000 | 0.810 | 0.714 |
| T1088 | 0.954 | 1.000 | 0.778 |
| T1089 | 0.972 | 0.813 | 0.624 |
| T1090 | 0.977 | 0.870 | 0.399 |
| T1091 | 0.832 | 0.571 | 0.071 |
| T1092 | 0.704 | 0.782 | 0.382 |
| T1093 | 0.673 | 0.519 | 0.109 |
| T1094 | 0.649 | 0.580 | 0.144 |
| T1095 | 0.722 | 0.711 | 0.448 |
| T1096 | 0.766 | 0.421 | 0.098 |
| T1099 | 0.800 | 0.375 | 0.101 |
| T1100 | 0.883 | 0.820 | 0.258 |
| T1101 | 0.960 | 0.988 | 0.783 |

We also investigated the impact of alignment depth and sequence length on contact prediction on the CASP14 dataset. For this we segmented the targets into groups with either less than 400 sequences, or between 400 and 15,000 sequences (threshold of maximum MSA depth). Figure 2 shows that a correlation between shallow MSAs and average prediction accuracy exists with a person correlation of *r > 0*.*7*. However, for deeper MSAs this correlation is no longer observed. Further, we compared the dependency of prediction accuracy on the sequence length of the target and found no correlation with *r = 0*. From this, we conclude sequence length independence of ProSPr and that finding at least a few hundred sequences is helpful to increase the predictive performance of ProSPr, but deeper alignments hold no clear benefit.

**Figure 2.**
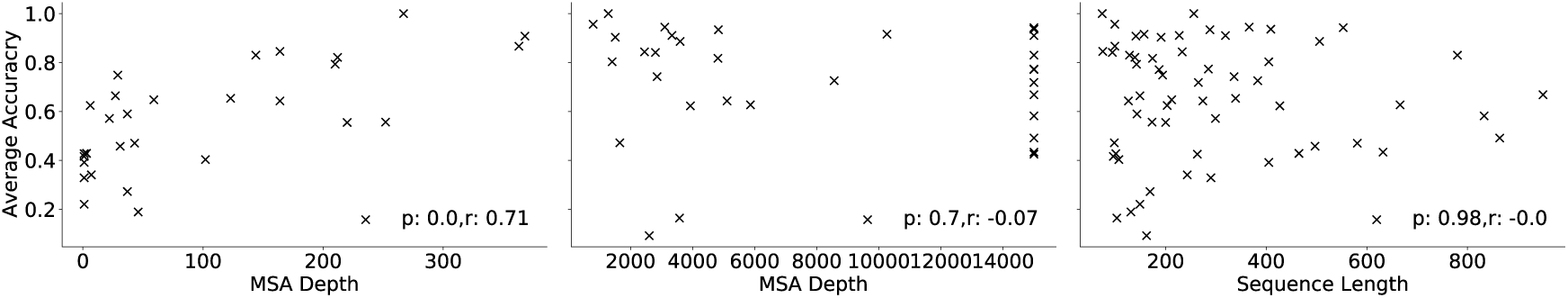
On left correlation analysis of average accuracy for CASP14 targets with MSA smaller than 400 sequences. Middle, correlation analysis for MSA deeper than 400 sequences. On right, correlation analysis of average accuracy and target amino acid sequence length.

**Figure 3.**
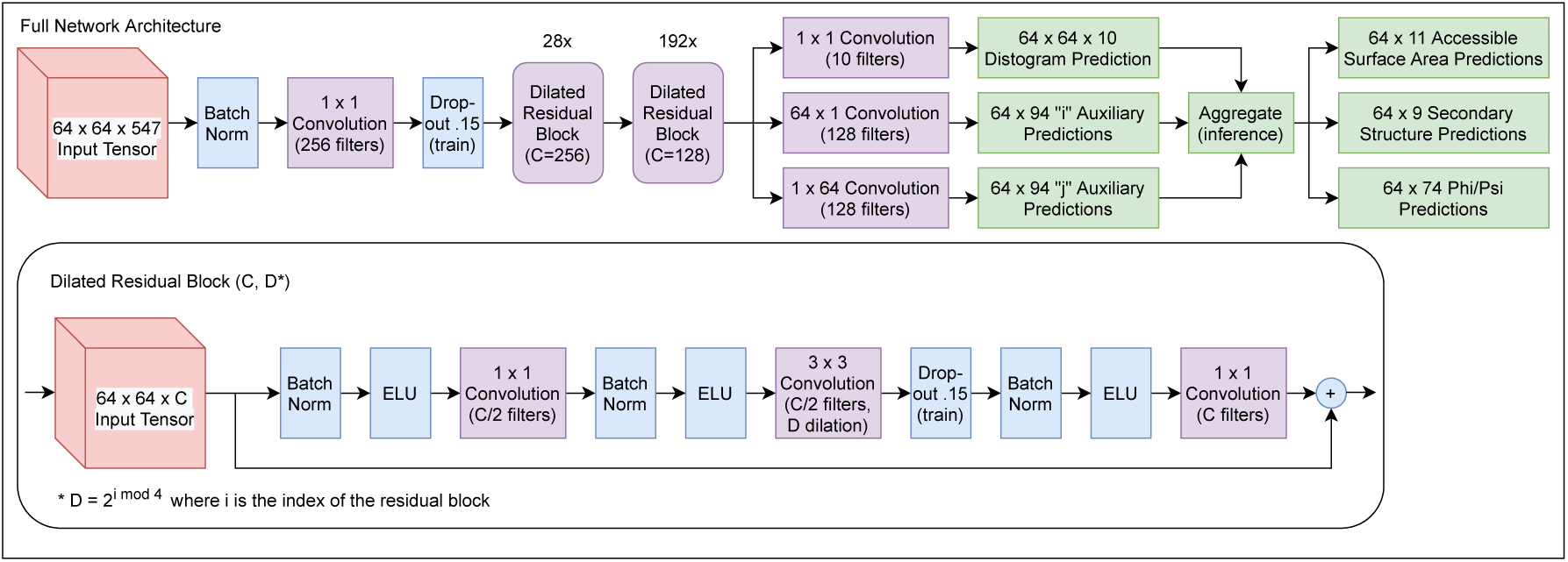
ProSPr Network Architecture

**Figure 4.**
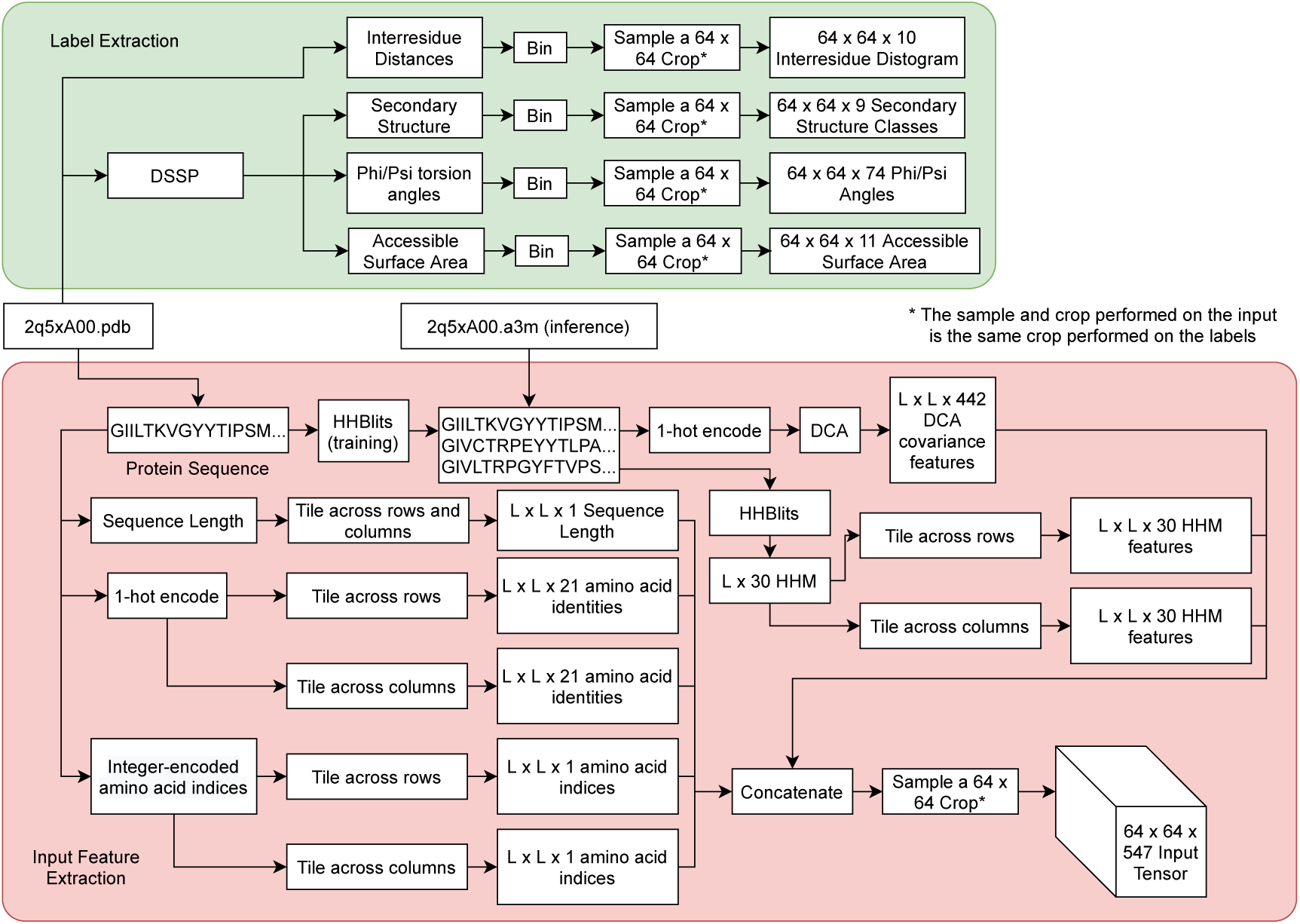
Detailed view of ProSPr data pipeline. For training a protein structure in pdb file format is used to create inputs and labels. For inference, a multiple sequence alignment in a3m file format is expected.

## 3. Methods

### ProSPr Overview

ProSPr predicts a series of features related to three-dimensional protein structures that can be referred to as protein structure annotations [16] (PSAs). The primary purpose of ProSPr is to predict the distances between pairs of residues for a given sequence. Specifically, this is defined as the distance between the Cβ atoms of two residues *i* and *j* (Cα is used in the case of glycine). ProSPr also predicts secondary structure (SS) classes, relative accessible surface area (ASA), and torsion angles for each residue in a sequence. However, these are included only as auxiliary features to improve the quality of the distance predictions (see Methods).

All ProSPr predictions are categorical in nature, and otherwise continuous values have been discretized into bins. For example, the inter-residue distances were divided into 10 bins: < 4 Å, 4 ≤ d < 6 Å, 6 ≤ d < 6 Å, … etc. until the final bin which included all distances greater than or equal to 20 Å. This specific format was developed in alignment with the distance prediction format announced for CASP14 [17].

#### Model Architecture

ProSPr is a deep, two-dimensional convolutional residual neural network[18] whose architecture was inspired by that of the 2018 version of AlphaFold1[8]. After performing an initial BatchNorm [19] and 1×1 convolution on the input tensor, the result is fed through the 220 dilated residual blocks that make up the bulk of the network. Each block consists of a BatchNorm followed by an exponential linear unit (ELU) activation[20] and a 1×1 convolution, then another BatchNorm and ELU, a 3×3 dilated convolution[21], and finally another BatchNorm, ELU, a 1×1 projection, and an identity addition. The blocks cycle through 3×3 convolutions with dilation factors of 1, 2, 4, and 8. The first 28 of these blocks use 256 channels, but the last 192 only use 128. Once passed through all 220 blocks, a 1×1 convolution is applied to change the number of channels down to 10 for distance predictions, while 64×1 and 1×64 convolutions are applied to extract the *i* and *j* auxiliary predictions, respectively.

### Input Features

The input tensor to ProSPr has dimensions *L x L x 547* and contains both sequence and MSA-derived features. The sequence information is provided as 21 one-hot encoded values, 20 for the natural amino acids and another for unnatural residues, gaps, or padding. The residue index information is also included as integer values relative to the start of the sequence. A Hidden Markov Model (HHM) is constructed from the MSA using HHBlits [22], for which numerical values are directly encoded as layers in the input tensor. Finally, 442 layers come from a custom direct-coupling analysis[23] (DCA) computed from the raw MSA.[24] Further details can be found in the released code, which includes a function for constructing full input from sequence and MSA.

### Training Data

We derived the data used to train these ProSPr models from the structures of protein domains in the CATH s35 dataset[25]. First, the sequences were extracted from the structure files. We then constructed multiple sequence alignments (MSAs) for each sequence using HHBlits[22] (E-value 0.001, 3 iterations, limit 15,000 sequences). Inter-residue distance labels were calculated from the CATH structure files and binned into 10 possible values, in accordance with CASP14 formatting as described previously. We then used the DSSP algorithm[26] to extract labels for secondary structure (9 classes native to DSSP), torsion angles (phi and psi, each sorted into 36 10º bins from -180º to 180º, plus one for error/gap) and relative accessible surface area (ASA) (divided into 10 equal bins, plus another for N/A or gap).

### Training Strategy

After generating the input data and labels for the CATH s35 domains, we split them into training (27,330 domains) and validation sets (2,067 domains). To augment the effective training set size, we used two strategies. First, we constructed ProSPr so that it predicts 64×64 residue crops of the final distance map. By doing this, we transformed ∼27k domains into over 3.4 million training crops. In each training epoch, we randomly applied a grid over every protein domain to divide it into a series of non-overlapping crops. Doing this each epoch also increased the variety of the input since the crops were unlikely to be in the same positions each time. Second, we randomly subsampled 50% of the MSA for each domain in each epoch. From this smaller MSA, we calculated the HHM and DCA features used in the input vector. This strategy also served to increase the variety of the training data used by the network to prevent overfitting.

All models were trained using a multicomponent cross-entropy loss function. The overall objective was to predict accurate inter-residue distances, the secondary structure (SS), torsion angles (phi/psi), and accessible surface area (ASA) tasks were included as auxiliary losses with the idea that adding components that require shared understanding with the main task could improve performance. Each of the cross-entropy losses was weighted by the following terms and summed to make up the overall loss: 0.5 SS, 0.25 phi, 0.25 psi, 0.5 ASA, and 15 for the distances.

All models used 15% dropout and an Adam optimizer with an initial learning rate (LR) of 0.001. The LR of model A decayed to 0.0005 at epoch 5 and further to 0.0001 at epoch 15. For model B the LR decreased to 0.0005 at epoch 10 and then to 0.0001 at epoch 25. Lastly, the LR of model C dropped to 0.0005 at epoch 8, and down to 0.0001 at epoch 20.

Each model trained on a single GPU (Nvidia Quadro RTX 5000 with 16GB) with a batch size of 8 for between 100 and 140 epochs, which took about two months. The validation set was used as an early-stopping criterion (using static 64×64 crop grids to reduce noise) and the three checkpoints of each model with the lowest validation losses were selected for testing. The CASP13 test set was then used for final model selection, and the CASP14 predictions were made and analyzed as described earlier.

### Inference

At inference time, we take crops that guarantee coverage of the entire sequence and take additional random crops to cover boundaries between the original crops. We then predict all features for each crop and average the aggregated predictions. The aggregation step consists of aggregating predictions across all crops for each pair *i,j* of indices (in the case of distance predictions), and each index *i* (in the case of auxiliary predictions), then taking the average prediction across all crops. Due to this cropping scheme, some crops will aggregate more predictions than others, which is corrected for through averaging.

## 4. Conclusion

We developed an updated version of the ProSPr distance prediction network and trained three new models. We found that an ensemble of all three models yields the best performance on the CASP14 test set, which agrees with our previous finding that deep learning models are frequently complimentary. We further investigated the dependency on multiple sequence alignment depth and found that very shallow alignments reduce the accuracy of the network but adding more sequences beyond a few hundred to an alignment does not result in further performance gains. We found that contact prediction accuracies for ProSPr on the CASP14 dataset are of high quality for short and mid contacts but lacking for long contacts. This is likely due to our strategy used for creating multiple sequence alignments, which does not leverage genomic datasets and results frequently in very shallow alignments. We also found that amino acid sequence length does not correlate with contact prediction accuracy on the CASP14 test set.

Besides the technical discussion, this work also describes a comprehensive architecture of ProSPr and a training strategy, together with necessary scripts to enable rapid reproduction. To our knowledge, this is the first deep learning-based method for protein structure prediction that publishes not only models but reproducible training scripts. As such it might prove a very useful educational tool for students trying to understand the applications of deep learning in this rapidly evolving field.[27] The full training routine and necessary datasets are available to enable other groups to rapidly build on our models. All necessary tools and datasets can be found at https://github.com/dellacortelab/prospr.

## Author Contributions

DC and WB conceived this study. WB, JS, BH, DC trained the networks and performed the analysis. OF supported all other authors with the writing of the article.

## Data Availability Statement

All data is available on https://github.com/dellacortelab/prospr.

